# Human tau compromises neuronal structural integrity in *C. elegans* independently of disease-associated modifications

**DOI:** 10.1101/2025.08.01.668095

**Authors:** Maithili Joshi, Vera van Schijndel, Jorieke Tiggelaar, Sanne J.A. van Falier, Tessa Sinnige

## Abstract

Tauopathies are a group of progressive neurodegenerative diseases amongst which Alzheimer’s disease is the most prevalent. They typically have a late age of onset and lead to cognitive decline. Tau is a highly soluble, intrinsically disordered microtubule binding protein that aggregates in these diseases. The mechanisms by which age-associated changes alter tau functionality and compromise neuronal structural integrity remain poorly understood. Here, we show with the use of single-copy gene insertion *Caenorhabditis elegans* models that human tau preferentially localises to neuronal processes in the nematode, consistent with its function of binding to microtubules. The expression of tau in the nervous system led to age-associated changes in the structural integrity of the ventral nerve cord, in agreement with previous reports in touch receptor neurons. However, expressing either the disease-related P301L mutant, phosphomimic tau or co-expressing human kinases did not worsen the observed buckling phenotype. On the other hand, kinase expression was associated with the formation of tau inclusions in a small proportion of animals at old age. These results suggest that human tau induces structural alterations in *C. elegans* neurons independently of pathological phosphorylation or aggregation. At the same time, the late-onset formation of tau inclusions upon kinase overexpression make this new model suitable to study the sequence of events leading to tau pathology.

## Introduction

Tauopathies are a heterogenous group of neurodegenerative diseases characterised by the accumulation of the protein tau in the form of neurofibrillary tangles (Creekmore et al., 2024; Götz et al., 2019; Wang & Mandelkow, 2016). Primary tauopathies such as frontotemporal dementia (FTD) are driven by mutations in tau, and display neuronal and glial tau tangles as the main pathological hallmark. Alzheimer’s disease (AD) is the most prevalent secondary tauopathy, featuring extracellular amyloid plaques next to intraneuronal tau tangles. AD is largely sporadic, with age being the main risk factor next to genetic background and lifestyle (Scheltens et al., 2021).

Tau is an intrinsically disordered protein known to bind to microtubules. It is produced as six different isoforms, varying from 352 to 441 amino acids in length. These isoforms carry either 0, 1 or 2 N-terminal inserts combined with either 3 or 4 tandem repeats of an amino acid motif which is the attachment site to microtubules, known as the microtubule binding repeat region (MTBR) (Götz et al., 2019). In humans, tau is distributed throughout the neuron in early development, whereas in mature neurons it is localised to axons. Tau pathology is accompanied by mislocalisation to the cell body, where neurofibrillary tangles are found (Götz et al., 2019).

Post-translational modifications are known to play a key role in this pathological process, most notably phosphorylation (Alquezar et al., 2021; Basheer et al., 2023; Wegmann et al., 2021). It is hypothesized that hyperphosphorylation of tau causes the protein to detach from microtubules, leading to an increased concentration of free cytosolic tau. This can lead to aggregation of tau and neurotoxicity (Wang & Mandelkow, 2016; Wegmann et al., 2021). However, the sequence of these events and the exact cause of toxicity remain unclear.

The tau protein and tau-associated pathologies are not unique to humans (Edler et al., 2017; Oikawa et al., 2010). Microtubule-associated proteins (MAPs) of the tau/MAP2/MAP4 family are found throughout much of the animal kingdom (Dehmelt & Halpain, 2005). The nematode *Caenorhabditis elegans* has one homologue of tau/MAP like proteins known as protein with tau like repeats-1 (PTL-1), which localises to both axons and dendrites and provides structural integrity to the neurons (Goedert et al., 1996). It is mostly expressed in the mechanosensory touch receptor neurons, which contain unusual 15 protofilament microtubules, in larval and adult stages (Goedert et al., 1996; Taylor et al., 2021).

*C. elegans* has been employed as a model organism to study both functional and pathological aspects of PTL-1 and human tau. PTL-1 was found to be important for maintaining neuronal health during ageing, and either increasing or decreasing its levels leads to loss of structural integrity of the touch receptor neurons (Chew et al., 2013). PTL-1 is involved in retrograde transport (Tien et al., 2011), and plays a role in mechanosensation but without visibly modulating microtubule organisation (Gordon et al., 2008).

Studies aiming to understand human tau pathology have been mostly based on neuronal overexpression of different wild type and mutant tau constructs. These typically reported functional defects accompanied by damage to neuronal structures (Brandt et al., 2009; Fatouros et al., 2012; Kraemer et al., 2003; Miyasaka et al., 2005; Pir et al., 2016). In some cases, insoluble tau was found to accumulate (Aquino Nunez et al., 2022; Fatouros et al., 2012; Kraemer et al., 2003), suggesting that tau aggregation is linked to the observed phenotypes.

In a different approach, wild type tau was expressed from a single-copy gene insertion in the touch receptor neurons, which did not lead to functional or morphological impairments. However, mutations mimicking certain post-translational modifications affected neuronal structure and function, while causing mitochondrial fragmentation (Guha et al., 2020) and impairing mitolysosome trafficking (Guha et al., 2022). In another recent study, panneuronal single-copy tau strains also lacked observable phenotypes, whereas multicopy strains displayed motility impairments correlating with expression levels (Han et al., 2024).

Despite these advances, the mechanisms by which disease-associated changes alter tau functionality and compromise neuronal structural integrity remain poorly understood. Here, we created single-copy gene insertion *C. elegans* models expressing human wild type and mutant tau in the nervous system and found that the ventral nerve cord displayed structural abnormalities. However, this phenotype was not affected by a disease-associated mutation or by phosphorylation, suggesting that it is related to the functional role of tau to bind to microtubules rather than to its pathological state.

## Results and discussion

### Human tau is enriched in neuronal processes in *C. elegans*

*C. elegans* single-copy strains were made based on the pan-neuronal expression of wild type and P301L mutant tau using the *rgef-1* promotor region. P301L is a disease associated mutation involved in frontotemporal dementia. For both strains, the 383 amino acid long 0N4R isoform of tau was used (Fig.1A), N-terminally tagged with monomeric enhanced yellow fluorescent protein (mEYFP). A single-copy strain expressing mEYFP with the same promotor served as the control.

Confocal imaging revealed that the protein localisation of wild type and mutant tau differs from that of mEYFP (Fig.1B, C, D). The fluorescence intensity is uniformly distributed across cell bodies and neurites in mEYFP. Contrastingly, wild type and mutant tau appear depleted in cell bodies and show a strong intensity in the nerve ring and ventral nerve cord (Fig. 1C, D arrows and arrow heads, respectively). Quantification of the ratio of fluorescence intensity in neurites versus cell bodies was done for the three strains. The ratio for mEYFP was close to 1, in agreement with a homogeneous distribution of the protein. For both wild type and mutant tau this ratio was around 1.7, suggesting that tau is enriched in the axons and dendrites compared to neuronal cell bodies (Fig. 1E). This preferential localisation of tau to neuronal processes is consistent with its function of binding to microtubules.

**Fig. 1.**
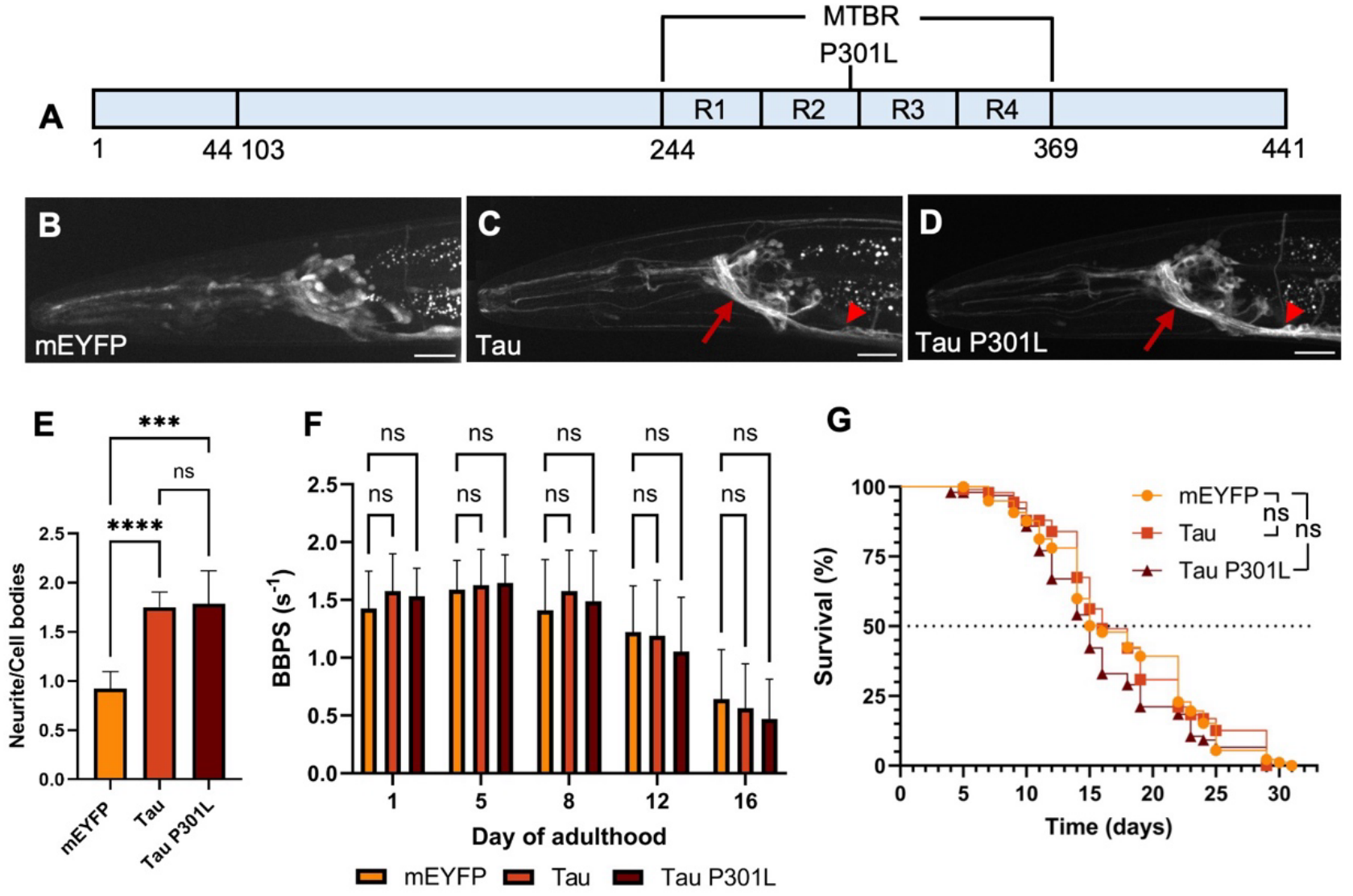
Human tau preferentially localises to neuronal processes in *C. elegans*. (A) Schematic design for tau 0N4R. Amino acids are numbered according to the longest tau isoform (2N4R). (B, C, D) Confocal images of the head region of mEYFP (B), tau (C) and tau P301L (D) *C. elegans* strains on day 1 of adulthood. Arrows show nerve ring; arrow heads the ventral nerve cord. Scale bars are 20 μm. (E) Quantification of the ratio of fluorescence intensities in neurites/cell bodies in mEYFP, tau and tau P301L strains on day 1 of adulthood (n = 8 animals per strain). Error bars are standard deviation. A Welch ANOVA was employed. (F) Quantification of the motility assay. n = 40 animals per condition (pooled from two biological repeats). Error bars are standard deviation. (For E and F, *p<0.05, ** p<0.01, ***p<0.001, ****p<0.0001, ns not significant.) (G) Survival assay. Dashed line indicates the median lifespan. n = 100 animals for each strain, representative of two biological repeats. Statistical analysis was done using a logrank test and the Gehan-Breslow-Wilcoxon test.

To assess the health of the animals upon the expression of tau, motility and survival assays were performed. As expected, we observed a reduction in motility at old age, but there were no significant differences between the three strains (Fig. 1F). Next, a survival assay was performed, which did not reveal significant differences between the strains either (Fig. 1G). Thus, we conclude that pan-neuronal single-copy expression of tau wild type or P301L is not considerably toxic in *C. elegans*.

### Tau expression causes buckling of the ventral nerve cord with age

To investigate age-related changes in the distribution of wild type and mutant tau, time course confocal imaging was done from day 1 to day 12 of adulthood. We found that some aged animals displayed a wavy appearance of the ventral nerve cord, which is a bundle of processes derived from interneurons and motor neurons (Fig. 2A). The wavy appearance has been previously reported to occur in ageing touch receptor neurons (Pan et al., 2011; Toth et al., 2012), and it has been linked to defects in β-spectrin (Krieg et al., 2014) as well as the microtubule cytoskeleton (Hsu et al., 2014; Krieg et al., 2017; Topalidou et al., 2012). This phenotype is also referred to as buckling, which is the term that we will use from here on.

**Fig. 2.**
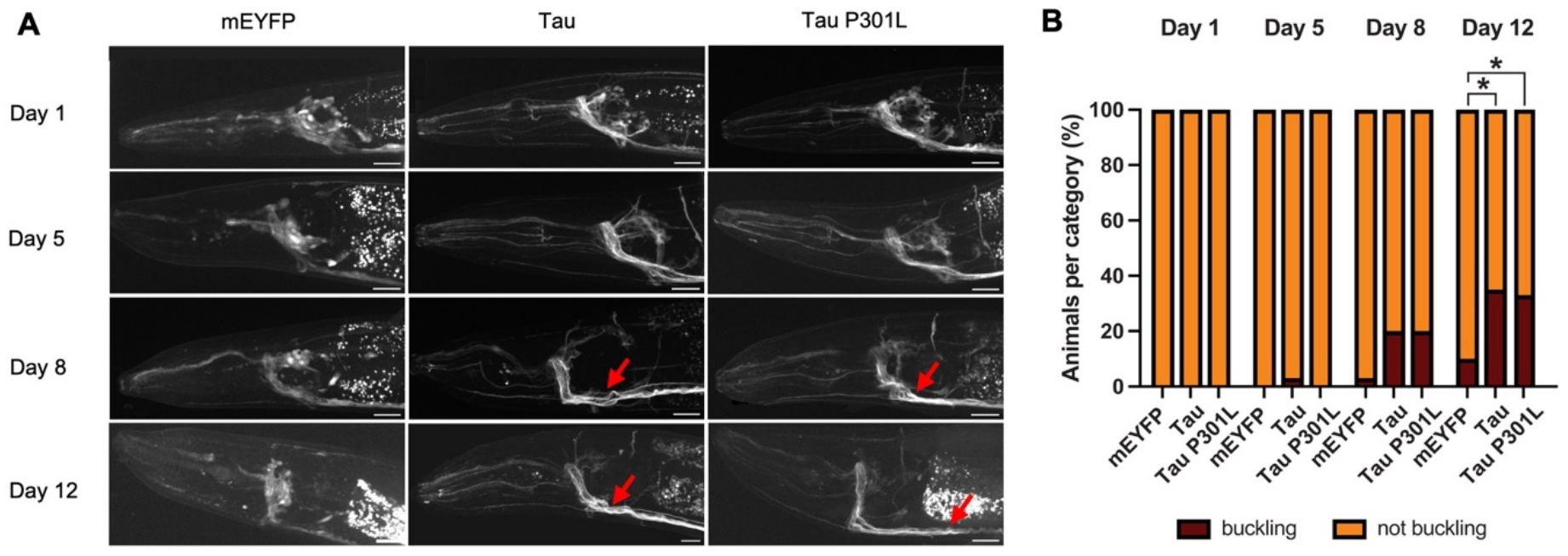
Tau expression induces buckling of the ventral nerve cord with age. (A) Representative confocal images of the head region of mEYFP, tau and tau P301L strains from day 1 to day 12 of adulthood. Buckling of the ventral nerve cord is indicated by red arrows. Scale bars are 20 μm. (B) Quantification of the percentage of animals showing buckling of the ventral nerve cord. n = 40 animals per condition pooled from two biological repeats. A Fisher’s exact test was employed for each strain and timepoint with respect to mEYFP (*p<0.05).

The percentage of animals displaying buckling of the ventral nerve cord increases with age, first appearing in the mEYFP control on day 8 (Fig. 2B). The presence of tau exacerbates this phenotype, appearing in a larger fraction of animals with statistical significance on day 12. Wild type and P301L tau show a comparable percentage of buckling animals at all timepoints (Fig. 2B), whereas they are expressed at similar levels, as validated from confocal images (Fig. S1). There results imply that the mutation does not worsen neuronal structural integrity compared to wild type tau. We did not observe other changes in protein distribution in the tau strains, such as macroscopically visible protein inclusions.

### Buckling phenotype is not affected by the tau phosphorylation

Tau hyperphosphorylation has been linked to its aggregation and the associated neurodegeneration. To understand the impact of tau phosphorylation on the structural integrity of the *C. elegans* nervous system, we created a single-copy model pan-neuronally expressing phosphomimic tau E14. This construct has 14 disease-associated serine/threonine phosphorylation sites mutated to glutamate to mimic phosphorylation (Fig. 3A), and has been reported to enhance toxicity in the brain of *Drosophila melanogaster* (Dias-Santagata et al., 2007; Khurana et al., 2006) and neuronal cell culture (Hallinan et al., 2019; Padmanabhan et al., 2024).

**Fig. 3.**
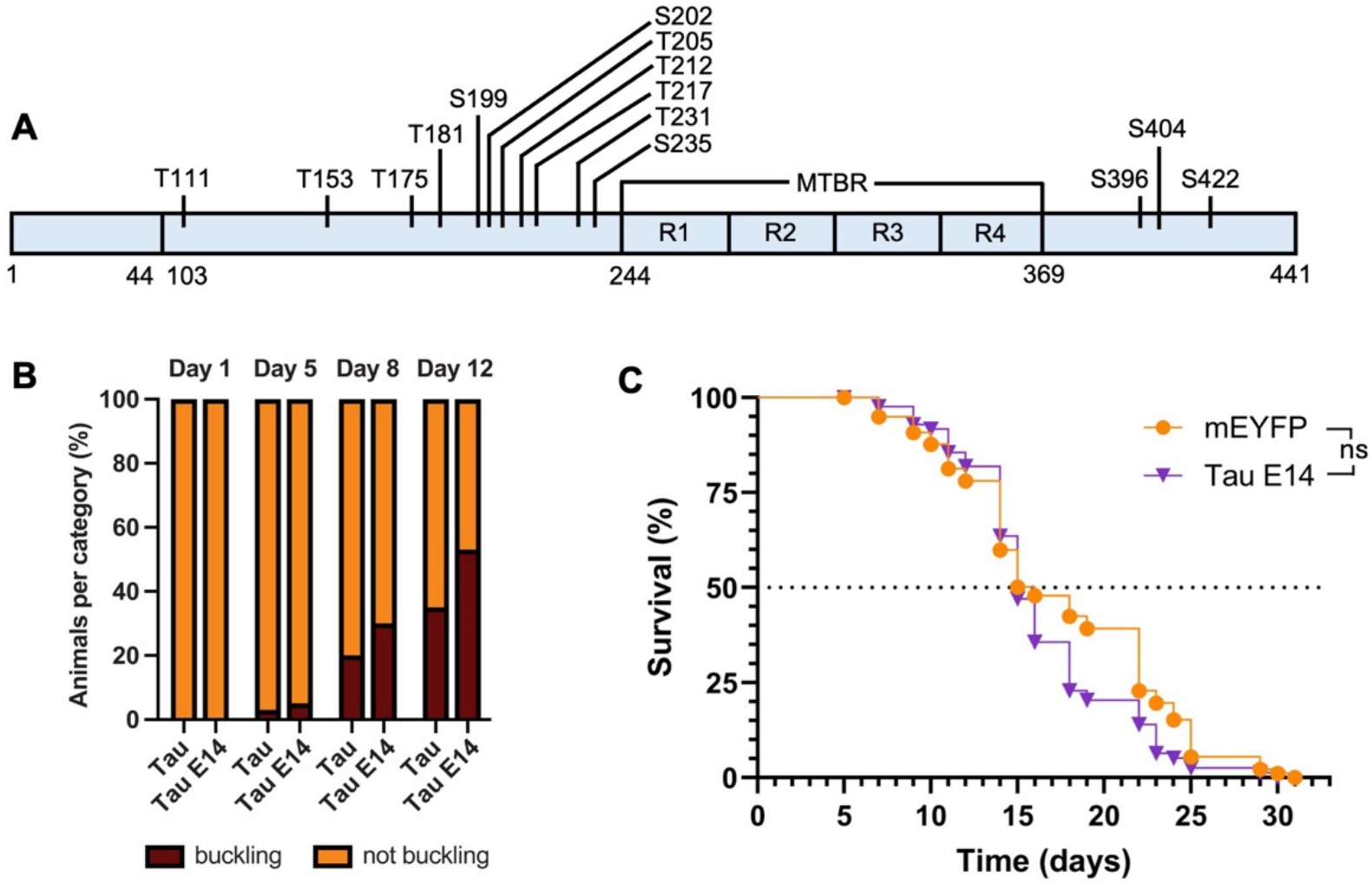
Expression of phosphomimic tau does not significantly increase tau-induced neuronal buckling. (A) Schematic showing the positions of the phosphomimic substitutions on the tau 0N4R construct. Amino acids are numbered according to the longest tau isoform (2N4R). (B) Quantification of the percentage of animals showing buckling of ventral nerve cord for tau, tau P301L and tau E14 strains. n = 40 animals per condition pooled from two biological repeats. A Fisher’s exact test was employed for each timepoint, resulting in no significant differences. (C) Quantification of the survival assay. n = 100 animals for each strain, representative of two biological repeats. Statistical analysis was done using a logrank test and the Gehan-Breslow-Wilcoxon test.

Expression of tau E14 resulted in a somewhat higher percentage of buckling compared to wild type tau, but this effect was not statistically significant (Fig. 3B). Thus, the phosphomimic substitutions do not appear to have a considerable impact on neuronal structural integrity in *C. elegans*. A survival assay was done to assess the health of these animals, and the lifespan of Tau E14 was not significantly different from mEYFP (Fig. 3C), indicating that expression of this pathogenic variant does not compromise overall organismal health.

Since phosphomimic substitutions have a different charge (-1) compared to phosphorylation (-2), they may impact the properties of the protein differentially. Hence, to also assess the effect of enzymatic phosphorylation, we overexpressed the human kinases GSK-3β and CDK5 pan-neuronally under control of the *rab-3* promotor in wild type animals. GSK-3β and CDK5 are both serine/threonine kinases that have been implicated in Alzheimer’s disease. CDK5 is reported to enhance both tau accumulation and neurodegeneration in *Drosophila* and mouse models (Noble et al., 2003; Saito et al., 2019). GSK-3β is involved in many cellular pathways and is another key kinase in tau hyperphosphorylation and neurodegeneration (Lauretti et al., 2020). Expression of GSK-3β or CDK5 in animals expressing wild type or P301L tau did not lead to significant changes in the percentage of buckling animals (Fig. 4A, B). The phenotype of controls expressing mEYFP was also largely unaffected by the kinases (Fig. S2).

**Fig. 4.**
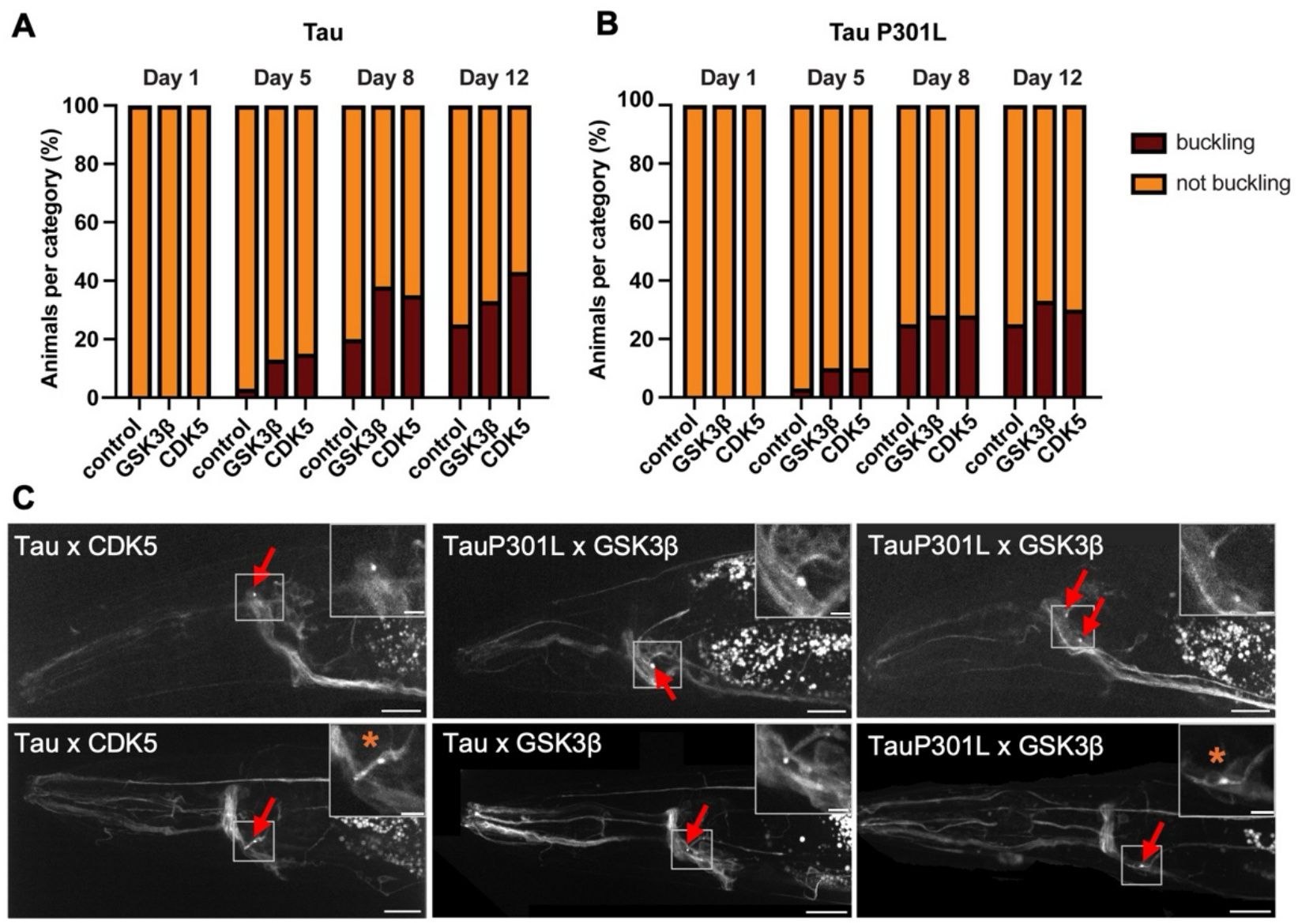
Expression of human kinases does not affect neuronal buckling, but leads to inclusion formation. (A) Quantification of the percentage of animals showing buckling of the ventral nerve cord with age for tau, tau x GSK-3β and tau x CDK5. (B) Quantification of the percentage of animals showing buckling of the ventral nerve cord with age for tau P301L, tau P301L x GSK-3β and tau P301L x CDK5. n = 40 animals per condition pooled from two biological repeats. A Fisher’s exact test was employed for each strain and timepoint with respect to control, resulting in no significant differences. (C) Confocal images of tau animals crossed with GSK-3β and CDK5 on day 10 (top row) and day 16 (bottom row) of adulthood. Foci are marked with red arrows. Scale bars are 20 μm. Insets show close-ups of the regions with foci, asterisks indicate those that appear perinuclear. Scale bars are 5 μm.

Confocal imaging of the strains expressing kinases on day 10 and day 16 of adulthood showed that some animals displayed tau foci, which appear as bright spots and are mostly localised to the posterior of the nerve ring rather than the ventral nerve cord (Fig. 4C). This was a rare occurrence with very few of the kinase expressing animals showing foci, yet none of the control animals displayed this feature as late as day 16 (Fig. S3). In some images, we could discriminate that the foci appeared to be localised to the cell body in a perinuclear region (asterisks in Fig. 4C). This observation strikingly resembles the formation of tau tangles in neuronal cell bodies, away from the physiological microtubule binding sites, that is observed in human Alzheimer’s disease.

In conclusion, we show that pan-neuronal single copy expression of human tau in *C. elegans* induces structural abberations of the ventral nerve cord, in agreement with previous observations in touch receptor neurons (Chew et al., 2013). However, this phenotype is not worsened by the disease-associated P301L mutation, nor by pathological phosphorylation states. These observations suggest that tau alters the cytoskeleton as a consequence of its physiological function to bind to microtubules. Interestingly, we observe late-onset inclusion formation upon overexpression of human kinases, providing an avenue to dissect the sequence of events leading up to tau pathology in future work.

## Materials and methods

### Molecular biology

Plasmids used in the study were generated using Gibson assembly (NEB) and are listed in Table S1. The pCFJ150 vector was used as backbone. For pan-neuronal expression, the promotor region of *rgef-1* was used. The promotor and 3’-UTR of *let-858* were amplified from *C. elegans* genomic DNA. EYFP was amplified from pPD30.38, which was a gift from Andrew Fire (Addgene plasmid # 1443; http://n2t.net/addgene:1443 ; RRID:Addgene_1443) and mutated (A206K) by site directed mutagenesis (Q5 site-directed mutagenesis kit, NEB) to create mEYFP. Tau was ordered as synthetic DNA from IDT. Site-directed mutagenesis was used to make Tau P301L (Phusion Site-Directed Mutagenesis Kit, Thermo Fisher Scientific). Tau E14, GSK-3β and CDK5 were ordered as synthetic DNA from Thermo Fisher Scientific. All oligonucleotides used in this study were ordered from IDT and are listed in Table S2.

### *C. elegans* strain generation

*C. elegans* strains used in this study are listed in Table S3. Single-copy strains were generated using the MosSCI method (Frøkjær-Jensen et al., 2008). An injection mix containing 10 ng/µL plasmid of interest, 50 ng/µL pCFJ601 (transposase), 10 ng/µL pMA122 (negative selection marker) and 30 ng/µL ccGFP (coelomocyte co-injection marker) was injected into the gonads of EG6699 adults. These were incubated at 25 °C for one week followed by a 2 h heat shock at 34 °C. The plates were chunked and screened for fluorescent animals with rescued movement and lacking expression of the co-injection maker. The insertion of the transgene was confirmed by amplifying the insert using PCR (primers NM3880 and NM3888) with LongAmp Taq DNA polymerase (NEB) followed by sequencing. The animals were backcrossed with N2 five times to dilute mutations induced by the heat shock. For the kinases, a mixture of 15 ng/µL of the plasmid of interest, 15 ng/µL of ccRFP and 70 ng/ µL 1 kb DNA ladder (NEB) was injected into N2 animals. The extra-chromosomal lines were then crossed with mEYFP, tau and tau P301L single copy lines.

### *C. elegans* strain maintenance

All strains were maintained on nematode growth media (NGM) seeded with *E. coli* OP50 at 15 °C. Age-synchronisation for time-course experiments was done by allowing adult animals to lay eggs for 2 h at 20 °C, followed by incubation at 20 °C for 3 days to reach day 1 of adulthood. All incubation steps and time course experiments were done at 20 °C, unless stated otherwise. Adult animals were transferred daily to fresh NGM plates to separate them from their offspring until they ceased to lay eggs.

### Microscopy

Leica MZ10F and Leica M165FC stereomicroscopes were used for the screening and selection of transgenic animals with filter set ET YFP with excitation 500/20 nm and emission 535/30 nm. Samples for confocal imaging or for scoring of neuronal buckling phenotype were prepared by immobilising adult animals on 2.5 % agarose pads in a drop of 10 mM tetramisole in M9 (22 mM KH2PO4, 42 mM Na2HPO4, 8.5 mM NaCl, 18.7 mM NH4Cl, 1 mM MgSO4). Confocal microscopy was done on a Nikon Eclipse Ti microscope body equipped with CSU-X1-A1 spinning disk (Yokogawa) and Nikon Plan Apo VC 60x /1.40 oil objective. The excitation and emissions wavelengths used for mEYFP were 488 nm and 525/50 nm respectively. The laser intensity used was 30 % and exposure time was 250 ms.

Quantification of fluorescence intensity in neurites and cell bodies was done by selecting regions of interest (ROI) in the nerve cord and the region posterior of the nerve ring that is enriched in cell bodies, respectively, in z-projected images. The integrated density and area values were obtained using image J and divided to calculate the average pixel intensities. Background subtraction was done using a ROI outside of the animal, and the average pixel intensities were divided to obtain the ratio of fluorescence in the neurites versus cell bodies.

To score animals for the buckling phenotype, a Leica DMi8 microscope with a Leica HC PL fluotar 40x oil objective with excitation wavelength 500/20 nm and emission wavelength 535/30 nm was used.

### Motility assays

For motility assays, 5-6 age synchronised animals were transferred to a drop of 75 µL M9 buffer in a 12 well plate, allowed to acclimatize for 30 s and then recorded for 30 s on a Leica S9i microscope at 26 fps. Animals were monitored until day 16 for the time course experiments, and on day 5 for the stress experiments (24 h post exposure to stress). A total of 40 animals per condition were analysed.

### Survival assays

100 age-synchronised animals were picked for each strain on day 1 of adulthood and grown at 20 °C. Animals were inspected daily and scored as dead if they were unresponsive to touch. This was done until all the animals were dead. Animals that were lost due to burrowing under the agar were censored and taken out of the analysis.

### Data analysis

All images and videos for motility assays were analysed using FIJI/ImageJ. The wrMTrck plugin in FIJI/ImageJ (Nussbaum-Krammer et al., 2015) was used to quantify body bends per second for motility assays. Statistical analysis was performed in GraphPad Prism 10. Motility assays were analysed using a two-way ANOVA. The comparison of the neurite/cell bodies ratios was analysed using a Brown-Forsythe and Welch ANOVA test. Analysis of lifespan was done using the Kaplan-Meier method with the logrank test and the Gehan-Breslow-Wilcoxon test. Statistical analysis of the buckling phenotype was performed using a Fisher’s exact test on 2x2 contingency tables, comparing the absolute numbers of buckling versus non-buckling animals for two strains in each test. The buckling graphs display the percentage of animals in each category. Motility data and fluorescent intensities are presented as mean values with standard deviation as error bars. Standard p values (*p<0.05, ** p<0.01, ***p<0.001, ****p<0.0001) were used for determining significance.

## Acknowledgements

We thank the Biology Imaging Centre at Utrecht University for the use of the confocal spinning disk microscope. We are grateful to Martin Harterink and Antoinette Killian for valuable discussions.

