## Supplementary information for "Human tau compromises neuronal structural integrity in *C. elegans* independently of disease-associated modifications"

**Table S1**: Plasmids used in this study

| Plasmid | Description |
|---|---|
| pCFJ150-pDEST [R4-R3] | Vector for single copy strains, has ampR |
| pDEST R3-R4 II | Overexpression vector, has ampR |
| pMJ20 | Pan-neuronal expression of mEYFP, back-bone: pCFJ150 |
| pMJ17 | Pan-neuronal expression of tau(WT), back-bone: pCFJ150 |
| pMJ18 | Pan-neuronal expression of tau(P301L), back-bone: pCFJ150 |
| pMJ22 | Pan-neuronal expression of CDK5, back-bone: pDEST R3-R4 II |
| pMJ23 | Pan-neuronal expression of GSK-3β, back-bone: pDEST R3-R4 II |
| pMJ24 | Pan-neuronal expression of tauE14, back-bone: pCFJ150 |
| ccRFP | RFP expression in coelomocytes, co-injection marker |
| ccGFP | GFP expression in coelomocytes, co-injection marker |

**Table S2**: Oligonucleotides used in this study

| Description | Sequence (5'-3') |
|---|---|
| p*rgef-1* forward primer | TAGAGGGTACCAGAGCTCACATCACAAATAAGGAATAAGCGACG |
| p*rgef-1* reverse primer | CATTTTTCTGAAAAGCCTGCTACGTACGAACGATTGAGCAGAAGG |
| mEYFP forward primer | AGGCTTTTCAGAAAAATGGACTACAAAGACGATGACGACAAG ATGGTGAGCAAGGGCGAG |
| mEYFP reverse primer | GCCCACTTTGTACAAGAAAGCTGGGTCCTTGTACAGCTCGTCCATGC |
| mEYFP position 2 reverse primer | GCCCACTTTGTACAAGAAAGCTGGGTCTTACTTGTACAGCTCGTCCATGC |
| Tau forward primer | GACCCAGCTTTCTTGTACAAAGTGGGCATGGCTGAGCCACGTCAAG |
| Tau reverse primer | GTCGATCATCCGATTCCACTTCACGATCGTTAGAGTCCTTGCTTGGCAAGG |
| *let-858* 3' UTR position 3 forward primer | GACCCAGCTTTCTTGTACAAAGTGGGCGATCGTGAAGTGGAATCGGATGA |
| *let-858* 3' UTR position 3 reverse primer | ACATATCCAGTCACTATGGCATACGGATTCGCATTTGCCAAG |
| *let-858* 3' UTR position 4 forward primer | CGATCGTGAAGTGGAATCGGATGA |
| *let-858* 3' UTR position 4 reverse primer | ACATATCCAGTCACTATGGCATACGGATTCGCATTTGCCAAG |
| p*rab-3* forward primer | AACATATCCAGTCACTATGGCATATTTTTTGACGACGACGAC |
| p*rab-3* reverse primer | CATTTTTCTGAAAAGCCTGCTACGTCTGAAAATAGGGCTACTGTAGATTTA |
| *let-858* reverse primer for overexpression | GAGAAAATACCGCATCAGGCATACGGATTCGCATTTGCCAAG |
| NM3880 | AGGCAGAATGTGAACAAGACTCG |
| NM3888 | ATCGGGAGGCGAACCTAACTG |

**Table S3**: *C. elegans* strains used in this study.

| Strain | Genotype | Available from |
|---|---|---|
| N2 | *C. elegans* wild isolate | CGC |
| EG6699 | *ttTi5605* II; *unc-119 (ed3)* III*; oxEx1578 [eft-3*p*::GFP* + *Cbr-unc-119 (+)]* | CGC |
| TSW58 (mEYFP single copy) | mbbSi6[*prgef-1*::FLAG::mEYFP::*let-858*-3'UTR + *Cbr-unc-119*] II; *unc-119(ed3)* III | This study |
| TSW59 (tau single copy) | mbbSi7[p*rgef-1*::FLAG::mEYFP:: Tau(WT)::*let-858*-3'UTR + *Cbr-unc-119*] II; *unc-119(ed3)* III | This study |
| TSW60 (tau P301L single copy) | mbbSi8[p*rgef-1*::FLAG::mEYFP:: Tau(P301L)::*let-858*-3'UTR + *Cbr-unc-119*] II; *unc-119(ed3)* III | This study |
| TSW47 (tau E14 single copy) | mbbSi11[p*rgef-1*::FLAG::mEYFP:: tauE14::*let-858*-3'UTR + *Cbr-unc-119*] II; *unc-119(ed3)* III | This study |
| TSW88 (CDK5) | mbbEx7[p*rab-3*::His::CDK-5::*let-858* 3'UTR + ccRFP] | This study |
| TSW89 (GSK3-β) | mbbEx8[p*rab-3*::His::GSK-β::*let-858* 3'UTR + ccRFP] | This study |
| TSW74 (cross of TSW58 and TSW88) | mbbSi6[p*rgef-1*::FLAG::mEYFP::*let-858*-3'UTR + *Cbr-unc-119*] II; *unc-119(ed3)* III; mbbEx7[p*rab-3*::His::CDK5::*let-858* 3'UTR + ccRFP] | This study |
| TSW75 (cross of TSW59 and TSW88) | mbbSi7[p*rgef-1*::FLAG::mEYFP:: Tau(WT)::*let-858*-3'UTR + *Cbr-unc-119*] II; *unc-119(ed3)* III; mbbEx7[p*rab-3*::His::CDK5::*let-858* 3'UTR + ccRFP] | This study |
| TSW76 (cross of TSW60 and TSW88) | mbbSi8[p*rgef-1*::FLAG::mEYFP:: Tau(P301L)::*let-858*-3'UTR + *Cbr-unc-119*] II; *unc-119(ed3)* III; mbbEx7[p*rab-3*::His::CDK5::*let-858* 3'UTR + ccRFP] | This study |
| TSW77 (cross of TSW58 and TSW89) | mbbSi6[p*rgef-1*::FLAG::mEYFP::*let-858*-3'UTR + *Cbr-unc-119*] II; *unc-119(ed3)* III; mbbEx8[p*rab-3*::His::GSK3-β::*let-858* 3'UTR + ccRFP] | This study |
| TSW78 (cross of TSW59 and TSW89) | mbbSi7[p*rgef-1*::FLAG::mEYFP:: Tau(WT)::*let-858*-3'UTR + *Cbr-unc-119*] II; *unc-119(ed3)* III; mbbEx8[p*rab-3*::His::GSK3-β::*let-858* 3'UTR + ccRFP] | This study |
| TSW79 (cross of TSW60 and TSW89) | mbbSi8[p*rgef-1*::FLAG::mEYFP:: Tau(P301L)::*let-858*-3'UTR + *Cbr-unc-119*] II; *unc-119(ed3)* III; mbbEx8[p*rab-3*::His::GSK-3β::*let-858* 3'UTR + ccRFP] | This study |

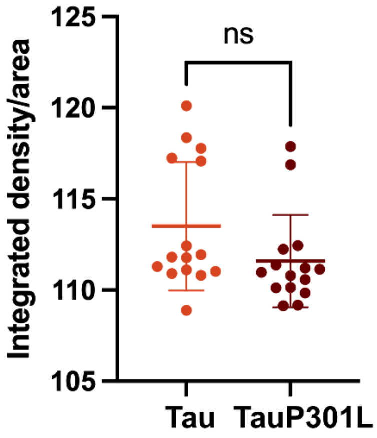


**Fig. S1** Quantification of integrated density/area of confocal images of day 1 animals. n = 15 animals per strain from one biological repeat. A t-test was performed (ns, not significant).

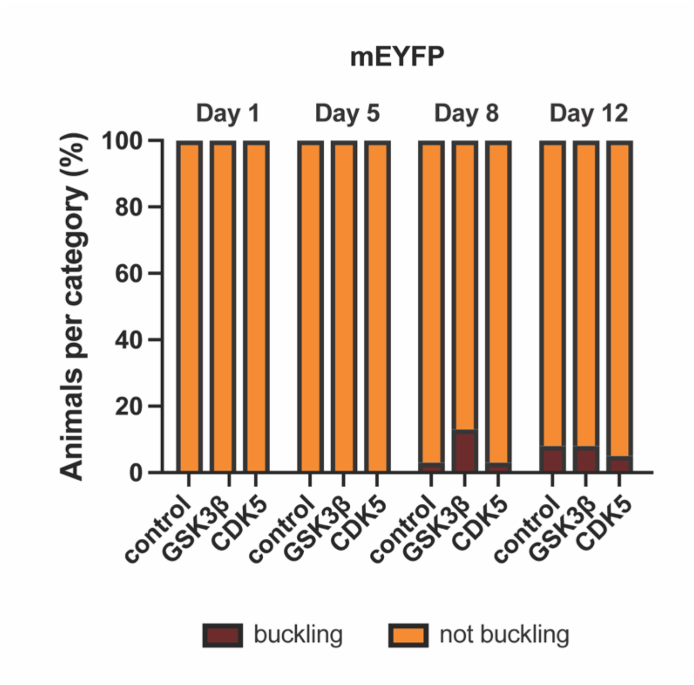


**Fig. S2** Quantification of the percentage of mEYFP animals crossed with GSK-3β and CDK5 showing neuronal buckling. n = 40 animals per condition pooled from 2 biological repeats. A Fisher's exact test was employed to assess the difference between mEYFP and each kinase at each timepoint, yielding no significant results.

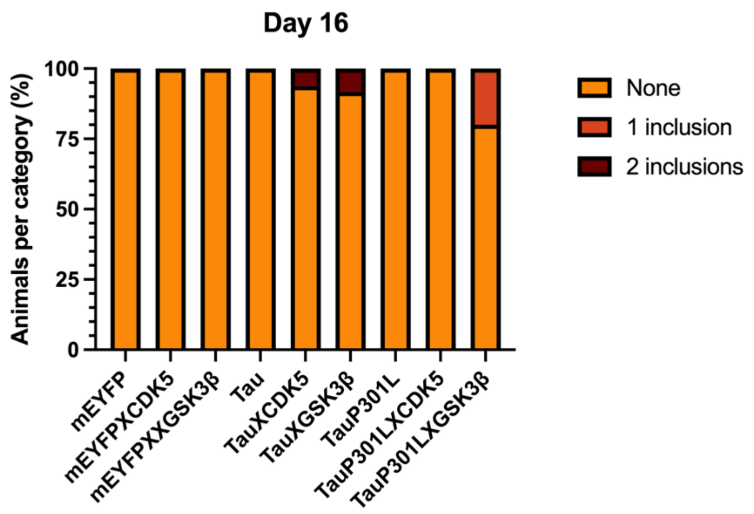


**Fig. S3** Quantification of inclusions seen on day 16 of adulthood in mEYFP, tau and tau P301L animals with and without expression of kinases. 5-20 animals were imaged per condition.
